# Cancer-associated DAXX mutations reveal a critical role for ATRX localization in ALT suppression

**DOI:** 10.1101/2024.11.18.624165

**Authors:** Sarah F. Clatterbuck Soper, Robert L. Walker, Marbin A. Pineda, Yuelin J. Zhu, James L. T. Dalgleish, Jasmine Wang, Paul S. Meltzer

## Abstract

To maintain genome stability, proliferating cells must enact a program of telomere maintenance. While most tumors maintain telomeres through the action of telomerase, a subset of tumors utilize a DNA-templated process termed Alternative Lengthening of Telomeres or ALT. ALT is associated with mutations in the ATRX/DAXX/H3.3 histone chaperone complex, which is responsible for deposition of non-replicative histone variant H3.3 at heterochromatic regions of the genome including telomeres. We wished to better understand the role DAXX plays in ALT suppression, and to determine which disease-associated DAXX mutations are unable to suppress ALT. To answer this question, we have leveraged the G292 cell line, in which ATRX is wild type but DAXX has undergone a fusion event with the non-canonical kinesin KIFC3. Restoration of wild-type DAXX in G292 localizes ATRX and abrogates ALT. Using this model system, we tested the ability of a panel of disease-associated DAXX missense variants to suppress ALT. Missense mutations in the ATRX binding domain, the histone binding domain, and the C-terminal SUMO interaction motif reduce the ability of DAXX to suppress ALT. Unexpectedly, we find that mutations in the DAXX histone binding domain lead to failure of ATRX localization. We conclude that a key function of DAXX in ALT suppression is the localization of ATRX to nuclear foci.

## Introduction

In proliferating cells, telomeres erode with each cell division due to the DNA end replication problem. This erosion is counteracted by telomerase, which adds telomere repeats utilizing an RNA template. In 10-15% of tumors telomeres are lengthened not by telomerase but through a DNA-templated process termed Alternative Lengthening of Telomeres or ALT^1^. ALT is more common in tumors of mesenchymal origin, including osteosarcoma, and is associated with mutations in the ATRX/DAXX histone H3.3 chaperone complex^2^. While ALT correlates strongly with *ATRX* mutations in osteosarcoma, *DAXX* mutations are frequently observed in pancreatic neuroendocrine tumors (PanNETs) and correlate with poorer prognosis^3–6^. Loss of ATRX or DAXX is often considered strong evidence of ALT, but it is less clear whether missense mutations in *DAXX* predispose cells to ALT telomere maintenance. We wished to understand more clearly which disease-associated missense mutations in *DAXX* are likely to confer an ALT phenotype, and how those mutations affect the function of DAXX.

DAXX has various cellular roles that could be implicated in ALT suppression. It was first described as an interactor of FAS and a modulator of apoptotic cell death^7,8^. DAXX is essential for embryonic development—knockout of Daxx in mouse embryos results in extensive apoptosis and embryonic lethality^9^. DAXX also co-regulates transcription of myriad target genes through its interactions with transcription factors and chromatin remodelers^10^. More recently, a role has been described for DAXX as a non-canonical protein folding chaperone^11^. In the context of ALT, DAXX is most associated with its activity as a chromatin remodeler. DAXX and the SWI/SNF helicase ATRX work in concert to deposit non-replicative histone variant H3.3 in heterochromatic regions of the genome, including telomeres^12,13^, and failure of H3.3 deposition is commonly thought to underpin ALT.

The structure of DAXX is largely disordered with two functional ordered domains. An N-terminal four-helix bundle (4HB) provides a platform for binding of ATRX and other interactors, while the histone binding domain enfolds the H3.3-H4 heterodimer^14–16^. DAXX ensures specificity for H3.3 by co-folding the with H3.3-H4 heterodimer and making contact with H3.3-specific residue G90^17^. The C-terminal region of DAXX is intrinsically disordered, and forms condensed droplets *in vitro*^18^. This region self-associates as well as interacting with binding partners such as SPOP^19^. Localization of DAXX within the nucleus is predominantly driven by its C-terminal SIM motif^20^. DAXX missense mutations in sarcomas with high ALT prevalence cluster in the H3.3 binding region of the protein, suggesting that H3.3 deposition may be key to ALT suppression^21^.

It is parsimonious to suppose that *ATRX/DAXX/H3.3* mutations potentiate ALT by reducing H3.3 deposition and thus altering chromatin states. Indeed, ALT is associated with reduced compaction of chromatin at telomeres^22^. To complicate this model, it has been found that H3.3 chaperone HIRA can compensate for H3.3 deposition at telomeres in ALT cells^23,24^. This result casts doubt on the model that H3.3 deposition by ATRX/DAXX is necessarily required for ALT suppression.

To determine which disease-associated *DAXX* mutations imply ALT, and dissect the function of DAXX in ALT suppression, we have leveraged the G292 osteosarcoma cell line. In G292 cells *ATRX* is wild type, but *DAXX* has undergone a fusion event with the non-canonical kinesin KIFC3^25,26^. Restoration of wild-type DAXX in G292 localizes ATRX and abrogates ALT. Using this model system, we tested the ability of a panel of cancer-associated *DAXX* missense variants to suppress ALT. We find that specific mutations throughout the DAXX protein, including both the DAXX H3.3 binding domain and the four-helix bundle, negatively impact the ability of DAXX to suppress ALT. Surprisingly, the ability of DAXX mutants to suppress ALT does not correlate well with H3.3 deposition at telomeres as measured by chromatin immunoprecipitation (ChIP). Instead, we find that the best correlate for the ability of DAXX to suppress ALT is colocalization of the DAXX variants with ATRX in nuclear foci. Importantly, most H3.3 binding mutants are also strongly impaired in ATRX localization. These results emphasize the importance of ATRX localization in ALT suppression and also represent the first direct characterization of the cellular defects of human-cancer-associated *DAXX* mutations.

## Results

### Creation of DAXX variant cell lines

To dissect the functional role of DAXX in ALT suppression, we selected a panel of single amino acid substitution mutants in DAXX. Utilizing the COSMIC database^21^, we chose nine missense mutations found in sarcomas and PanNETs spanning the length of the DAXX protein (Figure 1A). Of these, three (L130R, L134P, N153S) are located in or adjacent to the ATRX-DAXX binding interface^17^. Four of the sarcoma-associated mutations (S220P, N268S, A297P, G321D) are located in the DAXX histone binding domain. The Q469H mutation is located in the disordered DAXX acidic region. Finally, the D738V mutation alters the C-terminal SIM motif that is required for localization of DAXX at PML bodies^20^. This SIM motif is lost in the DAXX fusion protein found in G292^25,26^. We also tested two experimentally defined mutations known to alter the DAXX-H3.3 interface (F1A, noted in black). The Y222E mutation abrogates DAXX binding of H3.3, while E225A makes DAXX promiscuous for any H3 variant^15^.

**Figure 1.**
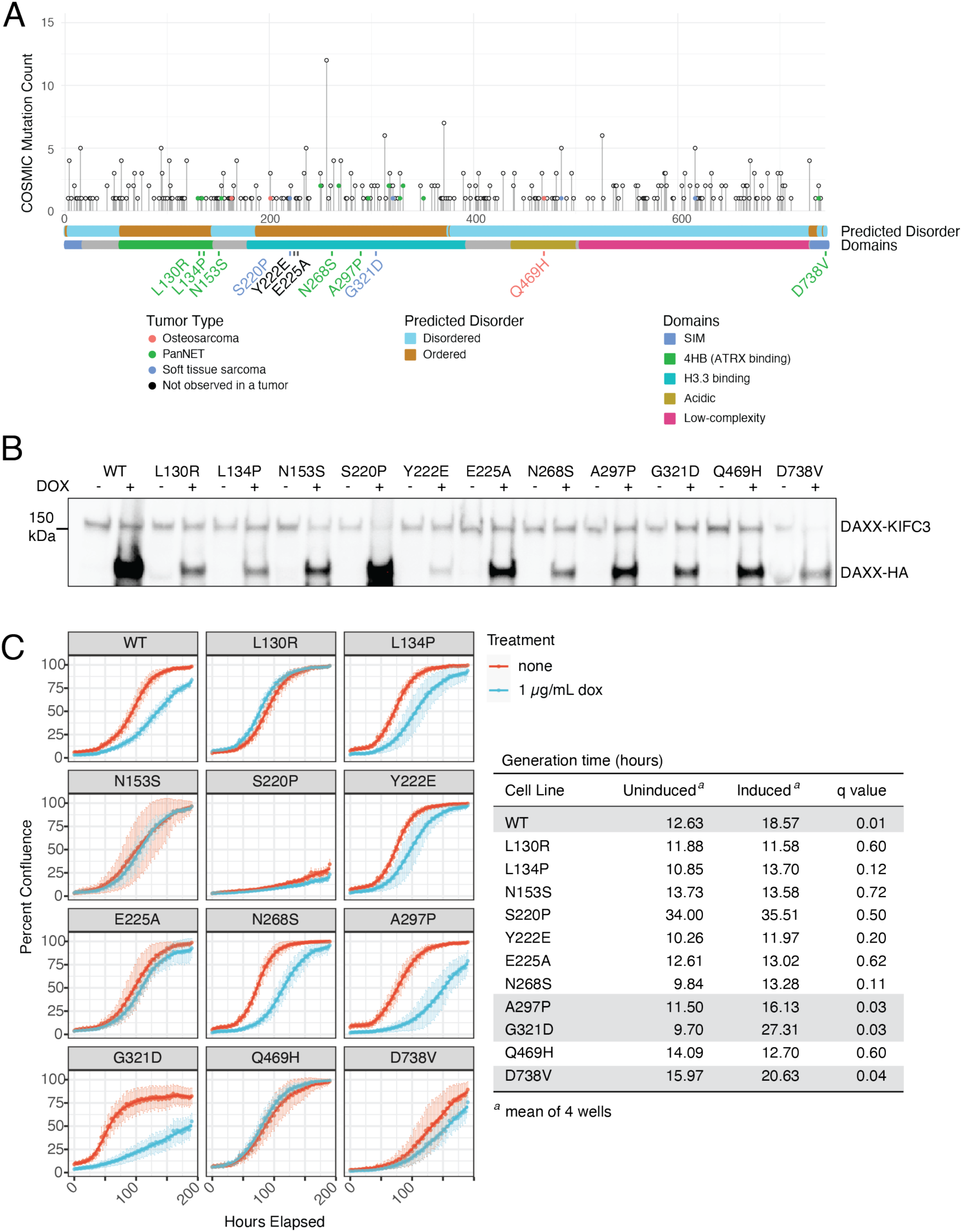
Selection of DAXX missense variants of interest and characterization of stable lines. A. Schematic of DAXX protein showing frequency of missense mutations as taking from the COSMIC database. Mutations found in non-epithelial tumors are colored by tumor type as described in the Tumor Type legend. Disordered domains and functional domains are marked as noted. Specific mutations selected for study are noted; text is colored for the tumor type in which that mutation was identified. Mutations noted in black were not observed in a tumor, but experimentally defined. B. Western blot showing DAXX expression with and without induction in cell lines from the selected mutants. DAXX was induced with 1µg/mL doxycycline. Expected molecular weights were 85 kDa for DAXX-HA and 126 kDa for DAXX-KIFC3 fusion. For most stable lines, DAXX-HA expression was higher than endogenous DAXX. C. Growth curves over the course of one week, with or without DAXX induction, taken from an Incucyte cell analyzer. Calculated generation times listed in the accompanying table reveal slowed cell growth with DAXX induction in several cases (highlighted in gray), including wild-type DAXX.

For accurate assessment of the ability of these mutants to suppress ALT, we used the pInducer-20 lentiviral vector to generate stable cell lines with inducible DAXX expression for each DAXX variant. Cell lines were analyzed as populations, except for DAXX-Y222E which was previously generated and cloned. Cell lines were continuously maintained under selection conditions to prevent silencing of the DAXX insertion locus. DAXX expression was induced with 1 µg/mL doxycycline (Figure 1B). In most cell lines this resulted in overexpression of the exogenous DAXX in comparison with endogenous protein, though DAXX-Y222E consistently yielded lower protein levels. This is consistent with prior observations that H3.3 binding stabilizes DAXX^17^.

To ensure that the variant DAXX proteins were not toxic, we calculated the generation time of each cell line with and without DAXX induction (Figure 1C). In some cases, including wild-type, DAXX induction resulted in a significantly slower generation time (see table). Consistent with prior FACS observations with wild-type DAXX induction^26^, every cell line continued to proliferate over the course of one week of DAXX induction. We noted that the S220P expressing line always grew more slowly. Based on nucleus size we believe this cell line underwent a genome duplication event during selection resulting in slow growth (Supplemental Figure 1).

### DAXX mutants form nuclear foci

Wild-type DAXX localizes predominantly to the nucleus, where it forms foci in addition to being distributed diffusely. Using immunofluorescence microscopy against the exogenous DAXX HA tag, we considered whether mutations affected the localization of DAXX (Figure 2A). We observed that all DAXX mutants localized predominantly to the nucleus, and all mutants localized to puncta to some extent. Using automated spot detection, we counted DAXX foci encompassing at least 16 pixels in each cell line. We observed that the L130R, L134P, and N153S mutants each formed significantly more foci per nucleus on average than wild-type DAXX (Figure 2B). Only the G321D mutant had a significantly decreased number of puncta. Calculating the average size of foci, we found that in the DAXX-L134P cell line the DAXX foci were significantly larger on average in addition to being more numerous than WT DAXX. However, foci of G321D and D738V variants were significantly smaller (Figure 2C). In the case of D738V this likely reflects the lack of PML body enrichment due to the impairment of the C-terminal SIM motif.

**Figure 2.**
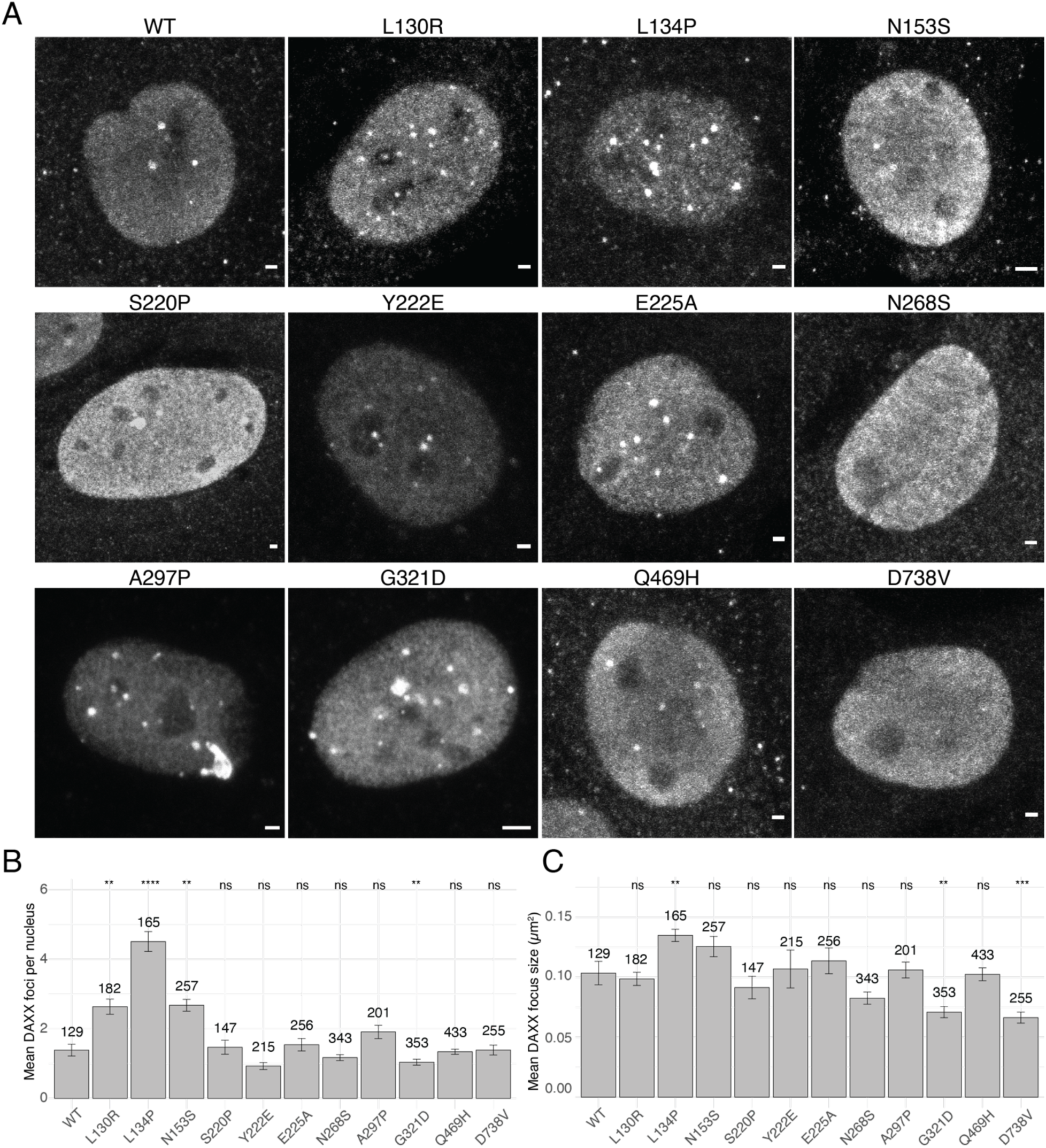
DAXX variants localize to nuclear foci. A. Immunofluorescence using an antibody against the HA tag to identify exogenous DAXX showed that all DAXX missense variants tested localized to nuclei. Representative nuclei are shown. Scale bars represent 2 µm. B. Enumeration of DAXX foci per nucleus. Foci of at least 16 pixels area were counted. Most DAXX variants formed about the same number of foci per nucleus, but mutations in the 4HB domain led to significantly more DAXX foci. Error bars represent s.e.m., number of nuclei analyzed is given for each cell line. Significance compared to wild-type DAXX by t-test, *: p ≤ 0.05, **: p ≤ 0.01, ****: p ≤ 0.0001 C. Average area of DAXX foci. While DAXX focus size was observed to be similar to wild type in most variants, DAXX-L134P foci were larger that wild-type DAXX in addition to being more numerous. The foci of DAXX variants G321D and D738V were smaller on average than wild-type DAXX. Error bars represent s.e.m., number of nuclei analyzed is given for each cell line. Significance compared to wild-type DAXX by t-test, *: p ≤ 0.05, **: p ≤ 0.01, ***: p ≤ 0.001

### DAXX HBD mutants poorly suppress ALT

Next, we assayed the capacity of each DAXX variant to suppress ALT. We relied on two measures of ALT activity to assess suppression: C-circles and “ALT-associated PML bodies” or APBs. C-circles are extra-chromosomal telomere repeats that are a hallmark of cells using ALT for telomere maintenance^27^. We tested C-circle levels after two days of DAXX induction and compared them to the uninduced cell lines (Figure 3A). As we have previously reported^26^, induction of wild-type DAXX expression in G292 significantly reduces C-circle production, reflecting suppression of the ALT phenotype. Of the panel of DAXX mutants, we found that induction of L130R, L134P, N153S, E225A, N268S, and Q469H significantly reduced detected C-circle levels, implying that these variants are at least moderately competent for ALT suppression. The DAXX variants S220P, Y222E, A297P, G321D and D738V were unable to suppress C-circle levels to a statistically significant extent. The S220P, Y222E, A297P and G321D mutations all fall within the histone H3.3 binding domain, implying a critical role for H3.3 binding in ALT suppression.

**Figure 3.**
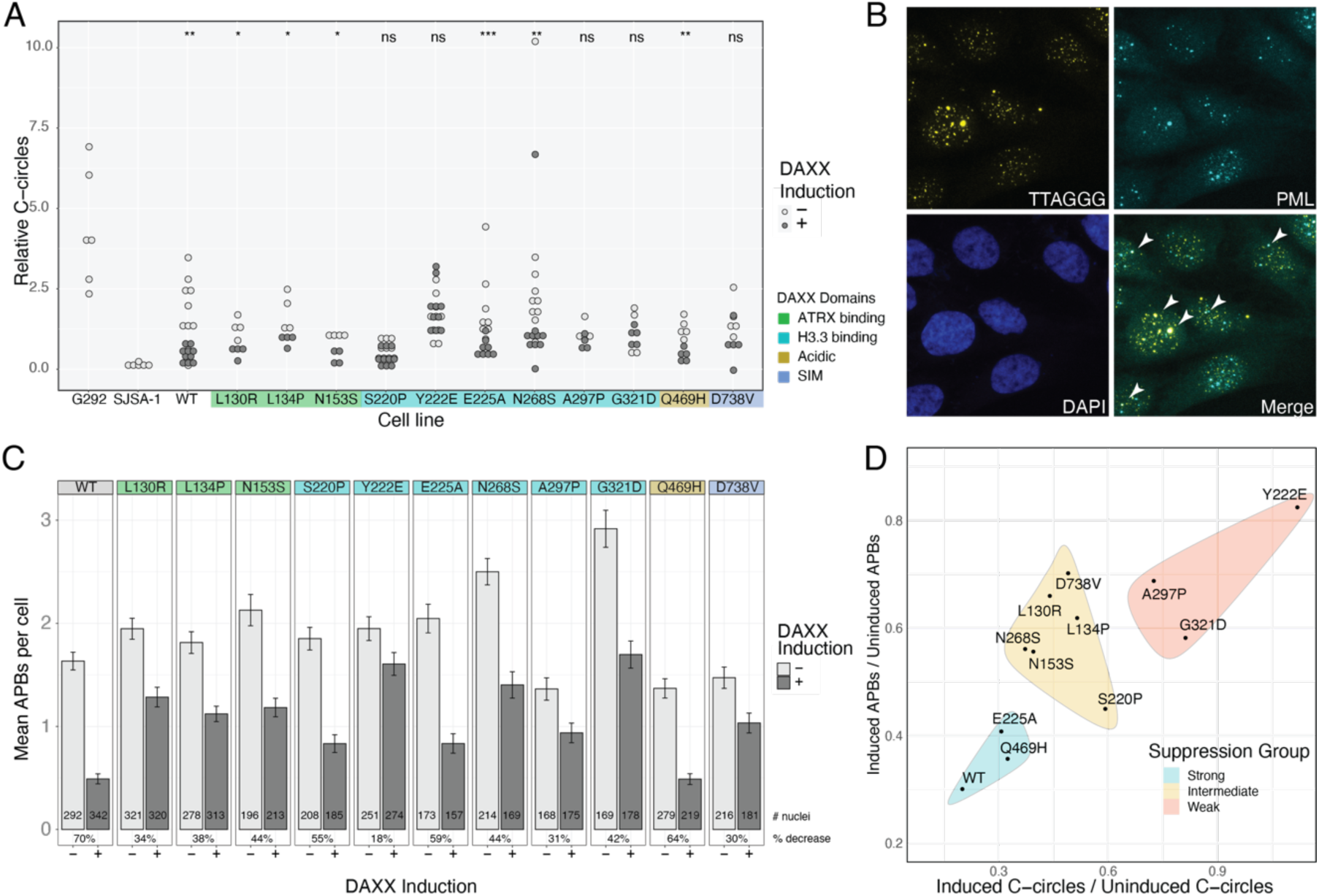
DAXX mutants vary in their capacity to suppress ALT. A. Comparison of relative C-circle levels in DAXX variants with and without DAXX induction. G292 parental and SJSA-1 represent ALT + and ALT – positive and negative controls, respectively. Each point represents a biological replicate. Statistical comparisons were made between uninduced and induced samples of the same cell line, using t-test, *: p ≤ 0.05, **: p ≤ 0.01, ***: p ≤ 0.001 B. Representative image of G292 cells showing IF-FISH to assay for APBs. Arrowheads point to APBs, where TTAGGG FISH signal overlaps with PML immunofluorescence. C. Enumeration of APBs per nucleus. Overlap areas of at least 4 pixels area were counted. DAXX induction decreased the number of APBs detected for all variants. Error bars represent s.e.m., number of nuclei analyzed is given for each cell line and treatment condition. Percent reduction of mean number of APBs with induction is provided for each cell line D. Scatterplot of ratio of C-circles with and without DAXX induction versus ratio of mean APBs per cell, with and without DAXX induction. DAXX mutants were categorized into groups conferring strong, intermediate or weak suppression based on reduction of C-circles.

In ALT, telomeres cluster at nuclear PML bodies forming APBs, where new telomere DNA is synthesized^28–30^. Along with C-circles, the presence of APBs is considered diagnostic of ALT^31^. To assay APBs, we performed telomere FISH in conjunction with immunofluorescence staining for PML as shown in Figure 3B, and counted overlapped foci programmatically. Surprisingly, we found that induction of DAXX significantly reduced the mean number of APBs per cell in every cell line (Figure 3C). Control experiments revealed that induction of NLS-mCherry from the pInducer vector also statistically decreased the mean number of APBs per cell, and induction of NLS-mCherry-3xSIM decreased APBs without affecting C-circles (Supplementary Figure 2). Previous reports have shown that ATRX, DAXX and PML are recruited to inducible CMV promoters in U2OS^32^, so we infer that the pervasive decrease in APBs is related to activation of expression from the inducible promoter, not a change in ALT status. Thus, we focused instead on the effect size of the decrease in APBs. We found that Y222E, A297P and D738V were least able to reduce the number of APBs per cell, while E225A and Q469H reduced APBs to an extent approaching WT DAXX. Other mutants exhibited an intermediate suppression phenotype.

Using reduction of C-circles as a guide, we categorized the DAXX mutants into three groups (Figure 3D). The Y222E, A297P and G321D mutants have very little capacity to suppress ALT. The L130R, L134P, N153S, S220P, N268S and D738V mutants have limited capacity to suppress ALT. Finally, the E225A and Q469H mutants suppress ALT nearly as well as WT DAXX. We noted that the three DAXX variants most compromised in the ability to suppress ALT have mutations in the H3.3 binding domain, including Y222E which has been shown experimentally to be defective in H3.3 binding^33^. These observations led us to more closely interrogate the contribution of H3.3 deposition to ALT suppression.

### H3.3 deposition at telomeres poorly correlates with ALT suppression

To isolate the contribution of H3.3 deposition at telomeres to the capacity to suppress ALT, we selected the DAXX mutants Y222E, E225A and N268S (from the weak, strong and intermediate ALT suppressing groups, respectively) to perform H3.3 chromatin immunoprecipitation (ChIP). To specifically determine changes in H3.3 deposition at telomeres, we pulled down chromatin using an antibody against histone H3.3, followed by blotting for telomere sequence (*e.g.,* Figure 4A). We compared cell lines with and without exogenous DAXX expression. We expected that wild-type DAXX expression would significantly increase H3.3 deposition at telomeres but found that the changes in the fraction of telomere sequence pulled down by H3.3 ChIP were modest, variable, and ultimately not statistically significant (Figure 4B). ChIP results from expression of the N268S mutant appeared similar to those of wild-type DAXX. As expected, expression of the H3.3 binding mutant Y222E did not increase H3.3 pulldown of telomere sequence, nor did the H3 promiscuous E225A mutant. Taken together, changes in H3.3 occupancy of telomeres with DAXX expression across these four cell lines did not correlate well with the capacity of the respective DAXX mutants to suppress ALT. While this assay would not be able to detect small changes in H3.3 occupancy of telomeres, these results persuaded us to consider other functions of DAXX as potentially more central to ALT suppression.

**Figure 4.**
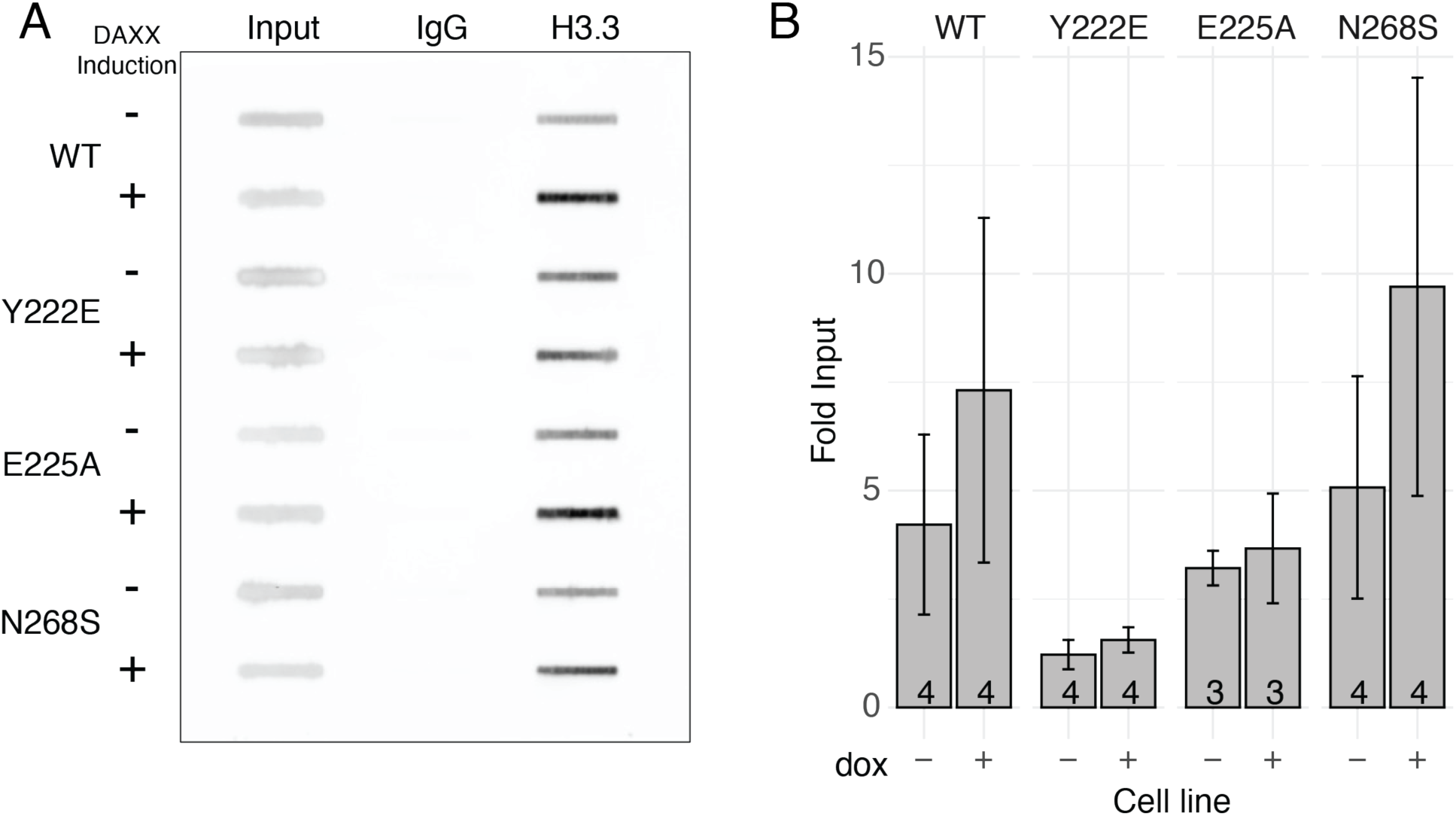
DAXX variant H3.3 deposition does not correlate with ALT suppression. A. Example blot of ChIP samples hybridized with probe against telomere repeats, showing telomere repeats bound by H3.3 in uninduced and induced conditions. B. Quantification of ChIP blots, normalized relative to input. Error bars represent s.e.m., number of biological replicates is noted on each bar.

### ATRX localization to PML bodies and telomeres is mediated by DAXX

The G292 cell line is WT for ATRX, and ATRX binds the DAXX-KIFC3 fusion protein^25,26^ In our previous studies, we observed that expression of WT DAXX in G292 led to colocalization of ATRX with DAXX and telomeres^26^. Consistent with this observation, we found that expression of WT DAXX increased ATRX localization to PML bodies (Figure 5A). To understand the functional importance of this localization, we performed ChIP-seq using an antibody against ATRX in G292 with and without WT DAXX expression. We found that expression of WT DAXX in G292 increased ATRX binding at telomeric repeats and at DNA sequences with >50% GC content (Figure 5B,C). This evidence supports a model in which DAXX mediates the correct localization of ATRX to telomeric sequences.

**Figure 5.**
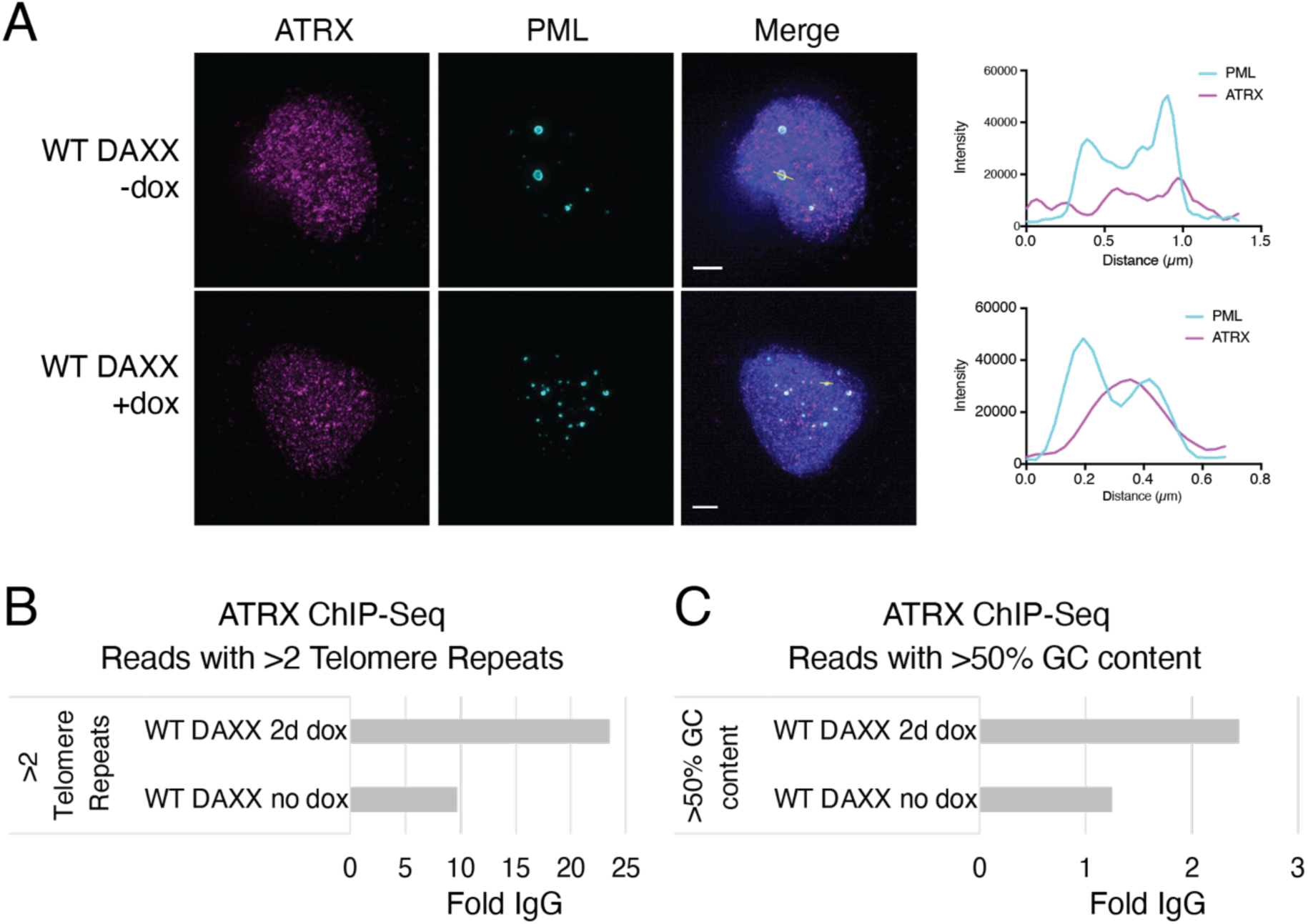
WT DAXX enhances localization of ATRX to PML foci and telomeric DNA sequences. A. Induction of wild-type DAXX expression results in stronger localization of ATRX to PML structures. Graphs represent intensity traces over the white line marking a PML focus in the corresponding image. Scale bars represent 2 µm. B. ATRX ChIP-seq shows enrichment of ATRX at telomere sequences with DAXX expression. Enrichment normalized to IgG controls C. ATRX ChIP-seq shows enrichment of ATRX at GC-rich sequences with DAXX expression. Enrichment normalized to IgG controls

### ATRX-DAXX foci correlate with ALT suppression

Given the importance of DAXX localization of ATRX to telomeres, we wished to understand the role of ATRX localization in ALT suppression. In G292, ATRX and endogenous mutant DAXX form complexes but do not localize to foci, and this failure to localize is permissive for ALT^25,26^. Thus, we expect the ability of DAXX to localize ATRX to nuclear foci is essential for ALT suppression. To assess the capacity of DAXX mutants to localize ATRX, we used confocal imaging to enumerate ATRX-DAXX colocalized foci (Figure 6A). In the G292 cell line, ATRX is predominantly diffuse, but becomes increasingly punctate with the expression of wild type DAXX (*e.g*., Figure 5A). A subset of ATRX and DAXX foci overlap, and these overlapped foci were enumerated on a per-cell basis (Figure 6B). We found that DAXX mutants E225A and Q469H, which exhibited robust ALT suppression, also had robust formation of ATRX-DAXX foci. Cells expressing DAXX-E225A were sometimes observed to have more than 10 ATRX/DAXX foci. We observed fewer ATRX-DAXX colocalized foci in cell lines expressing DAXX mutants with lower capacity for ALT suppression. Thus, we conclude that in most cases, the ability of DAXX to suppress ALT is tied to its capacity to localize ATRX to nuclear foci.

**Figure 6.**
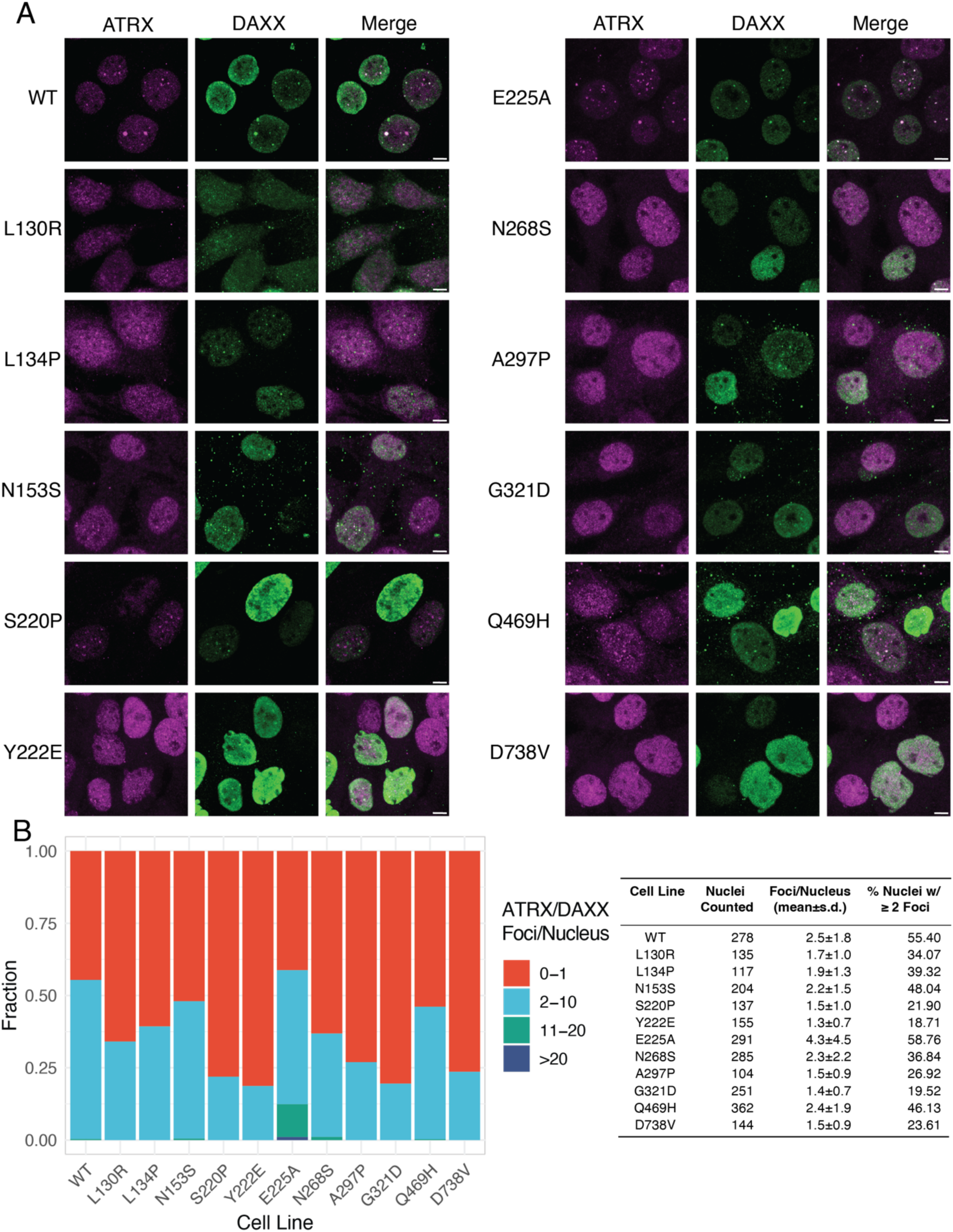
DAXX formation of foci with ATRX correlates with ALT suppression. A. Representative immunofluorescence images using antibodies against ATRX and against DAXX-HA. Scale bars represent 5 µm. B. Distribution of overlapped DAXX-HA and ATRX foci per nucleus in induced cells. Overlapped areas of at least 4 pixels area were counted. Most DAXX variants formed fewer ATRX overlapped foci on average than wild-type DAXX (see table). The E225A mutation led to significantly more ATRX-DAXX foci on average.

## Discussion

Here, we use a panel of disease-associated *DAXX* variants to address two intertwined questions. First, what is the essential function of DAXX in ALT suppression? And second, which disease-associated *DAXX* variants are permissive for ALT? We conclude that the essential requirement for ALT suppression is correct localization of ATRX within the nucleus as mediated by the DAXX C-terminal SIM motif. We observe that this localization can be disrupted by mutations in the 4HB domain, the histone binding-domain, or the C-terminal SIM motif. This implies that missense mutations throughout the DAXX protein sequence have the potential to lead to ALT.

In this work we have focused on DAXX in the context of an ALT+ osteosarcoma cell line that carries a DAXX fusion protein. Endogenous DAXX is expressed but fails to localize due to the loss of the C-terminal SIM motif. We are mindful of the fact that the DAXX variants we expressed would act in competition with endogenous DAXX for binding partners; for this reason, we overexpressed DAXX variants relative to endogenous DAXX. In G292, WT DAXX overexpression results in slower generation times, which may be attributable to the suppression of ALT and lack of telomere maintenance. None of the DAXX mutants we tested conferred a growth advantage in this cell line, regardless of their ability to suppress ALT, while several resulted in slower growth. Thus, we do not observe any gain-of-function growth advantages from the mutations we examined, at least in the context of an ALT cell line. Notably, the G292 cell line has a truncation in TP53, so we could not assess the proposed activity of DAXX in stabilizing or localizing p53^11,34^.

Consistent with the expectation that most DAXX mutations in ALT-prone cancers potentiate ALT, eight of the nine disease-associated mutations that we tested incompletely suppressed ALT. All the disease-associated DAXX variants had a stronger ALT-suppression phenotype than the experimentally defined Y222E mutant, however. In wild-type DAXX, Y222 forms part of a hydrophobic pocket that binds H3.3 residue I89. The Y222E mutation replaces the hydrophobic tyrosine residue with a charged glutamate, disrupting the ability of DAXX to bind H3.3^33^. In our hands the Y222E protein levels were consistently lower than those of any other DAXX variant, consistent with observations DAXX requires H3.3 binding for protein stability^17^. We speculate that low levels of DAXX-Y222E protein may drive the stronger phenotype we observed, and that such low DAXX protein expression may confer a selective disadvantage in the context of disease.

Of the disease-associated DAXX mutations that we tested, only the Q469H mutation, found in the disordered acidic region, suppressed ALT to an extent comparable to wild-type DAXX. A mutation in the disordered region of DAXX might be expected to affect the ability of the protein to form phase-separated foci, but Q469H formed foci at a comparable size and number to wild-type DAXX. Notably, this mutation was recorded in an osteosarcoma sample in which a DAXX E201D mutation in the histone binding region was also observed, though the zygosity of these mutations was not known^21^. It seems plausible, based on our observations, that the E201D DAXX mutation in this sample conferred an ALT phenotype, while the Q469H mutation acted as a neutral passenger.

It was our expectation that H3.3 deposition at telomeres would be essential for ALT suppression. We predicted that DAXX mutations that impaired H3.3 deposition at telomeres would be unable to suppress ALT. We were especially interested in the behavior of the E225A DAXX mutation, as it binds H3 histone variants promiscuously, and we judged it would serve as a test of whether H3.3 *per se* versus other H3 histones was essential for ALT suppression at telomeres. We found that DAXX E225A suppressed ALT to an extent comparable to wild-type DAXX, with no apparent deficits due to deposition of canonical H3. Using H3.3 ChIP and probing for telomere sequence, we found no correlation between extent of H3.3 deposition at telomeres and the capacity of wild-type, Y222E, E225A and N268S DAXX variants to suppress ALT. The ChIP experiment is an imperfect assay and would not detect small increases in H3.3 deposition with DAXX expression, but it led us to consider that H3.3 deposition might not be essential for ALT suppression by ATRX/DAXX. It has been demonstrated that HIRA can compensate for the ATRX/DAXX in depositing H3.3 at telomeric heterochromatin^23,24^, perhaps explaining the inconsistent increase in H3.3 we observe at telomeres with expression of wild-type DAXX. We conclude that H3.3 deposition at telomeres may not be directly required for ALT suppression.

Localization of telomeres to PML bodies is essential for the ALT mechanism^35^. Wild-type DAXX localizes to PML bodies via its C-terminal SIM motif^20^. In the G292 cell line, the DAXX-KIFC3 fusion protein has lost this SIM motif, resulting in a protein that binds ATRX but does not localize to PML bodies^25,26^. Thus, we find that to counteract the ALT mechanism that takes place at PML bodies, DAXX must both bind ATRX and correctly localize to PML bodies. We were satisfied to observe that the D738V SIM domain mutant, which is analogous to the DAXX-KIFC3 fusion endogenous to G292, poorly suppressed ALT, reinforcing the importance of the DAXX C-terminal SIM motif in ATRX/DAXX localization. We also observed that DAXX variants with mutations in the DAXX 4HB ATRX-binding region incompletely suppress ALT. We expect that the level of ALT suppression is likely related to the magnitude of the defect in ATRX binding; this may be a fruitful avenue for future exploration. Notably, DAXX variants with mutations in the ATRX binding site each were observed to form more and larger nuclear DAXX foci than other DAXX variants. It is possible that ATRX acts in competition with other DAXX binding partners, such as SPOP. In the absence of ATRX binding DAXX may form different complexes that are more prone to phase-separation. Alternatively, DAXX foci in the absence of ATRX may represent stalled biological assemblies that fail to resolve without ATRX activity.

Finally, we were most surprised to observe that DAXX variants with mutations in the histone binding domain fail to localize ATRX. We expected that these mutations would act as a separation of function, localizing ATRX but not depositing histone H3.3. This assumption has been made in other reports, where repletion of DAXX A297P was used as evidence for the importance of H3.3 deposition in ALT suppression^36^. We found instead that mutations throughout the histone binding domain negatively affect the localization of ATRX by DAXX. In the case of the Y222E mutation, it has been observed that the lack of H3.3 binding renders the variant unstable; this was consistently reflected in low protein expression in our hands^15^. It is possible that failure to localize ATRX is simply due to low protein levels in the Y222E expressing cell line. However, other DAXX histone binding variants expressed at levels significantly exceeding endogenous. Only one mutation (N268S DAXX) localized ATRX competently but did not fully suppress ALT. This variant merits further study to understand why it fails to suppress ALT completely.

To conclude, we find that mutations throughout the DAXX protein negatively impact the ability of DAXX to suppress ALT. We observe that ALT suppression is tied closely to ATRX localization, and that ATRX localization is negatively impacted by mutations in the H3.3 binding domain. We propose a model in which DAXX H3.3 binding is upstream of ATRX complex formation with the helicase role of ATRX at telomeres essential for ALT suppression.

## Methods

### Generation of DAXX mutant cell lines

G292 derived cell lines were maintained in McCoy’s 5A modified media supplemented with 15% fetal bovine serum and penicillin/streptomycin. The inducible WT DAXX cell line was generated as previously reported^26^. Briefly, wild-type DAXX cDNA was cloned into the pInducer20 expression vector (Addgene Plasmid #44012), lentivirus was generated using the psPAX and pMD2.G packaging vectors (Addgene Plasmid #12260 and #12259, respectively) and G292 cells were infected. Cells with successful insertions were selected with 500 µg/mL G418 and were subsequently maintained in 100 µg/mL G418. DAXX mutant expression vectors were derived from the WT DAXX expression vector using In-Fusion mutagenesis (Takara) with primers as listed in Supplementary Table 1. Stable cell lines were generated as described for the WT DAXX expression line. DAXX expression was induced with 1 µg/mL doxycycline for 48-72 hours before imaging and C-circle assays.

### Western blots

Western blots were performed as per standard procedures using nuclear extracts. To prepare nuclear extract, 2×10^6^ cells PBS-washed cell pellets were resuspended in 200 µL ice cold swelling buffer (10 mM HEPES pH 7.9, 10 mM KCl, 0.1 mM EDTA, 0.1 mM EGTA, 1mM DTT, 0.5 mM PMSF) and allowed to swell on ice for 15 minutes. Cell membranes were disrupted with 12.5 µL 10% NP-40 and nuclei were pelleted. Nuclei were resuspended in ice cold NTEN (20 mM Tris 8.0, 150 mM NaCl, 0.5 mM EDTA, 0.5% NP-40) and sonicated 3×5s on ice. Lysates were quantified using the standard BCA assay, denatured under reducing conditions in LDS buffer and 3% BME, and run on a polyacrylamide gel. Proteins were transferred onto PVDF membranes and detected with rabbit monoclonal anti-DAXX (Cell Signaling clone 25C12).

### Incucyte growth analysis

Cells were plated at a density of 10,000 per well in 96 well plates. After 24 hours, media was changed, and doxycycline was added to induction wells at 1 µg/mL concentration. Plates were transferred to the Incucyte S3 Cell Analysis device and imaging was initiated at this time. Images were taken at two-hour intervals for a minimum of one week and up to two weeks, and confluency was calculated for each time point. A minimum of four wells was imaged per condition. Generation times were derived from confluency data using Growthcurver^37^.

### C-circle quantification

C-circle levels were quantified using phi29 rolling circle amplification followed by dot blot. Genomic DNA was digested with 4 U/µg each HinfI and AluI, and 25 ng/µg RNase A for 2h at 37°C. For phi29 amplification, 16 ng of digested DNA was amplified with EquiPhi29™ DNA Polymerase (ThermoFisher) for 3h at 45°C. Minus-enzyme controls were run in parallel. Amplified and control reactions were blotted onto positively charged nylon membrane (Roche) and crosslinked. Membranes were probed with telomere specific 3′ labeled DIG probes (CCCTAA)^3^ (IDT) used at a concentration of 15 ng/mL. Detection was accomplished using the DIG Nucleic Acid Detection Kit (Roche). Images were quantified using ImageJ, and intensity values were normalized against the median intensity of each blot for comparison across experiments. A representative C-circle blot is shown in Supplementary Figure 3.

### Immunofluorescence

Cells were grown on acid-washed coverslips and were fixed with 4% paraformaldehyde for 15 minutes at room temperature. Following three washes with PBS, cells were permeabilized with 0.5% Triton X-100 for 5 minutes at room temperature, washed three times with PBS and blocked with 20% donkey serum for one hour at room temperature. Incubation with primary antibodies diluted in 5% donkey serum was carried out at 4°C overnight. Primary antibodies used included: rabbit monoclonal anti-ATRX (clone E5X70, Cell Signaling), rabbit monoclonal anti-DAXX (clone 25C12, Cell Signaling, 1:100 dilution), mouse monoclonal anti-PML (clone PG-M3, Santa Cruz, 1:200 dilution), mouse monoclonal anti-HA (clone 16B12, Biolegend, 1:100). Following primary antibody incubation, cells were washed three times with PBS and incubated with Alexa Fluor conjugated secondary antibodies (Jackson ImmunoResearch or ThermoFisher, 1:400 dilution) diluted in 5% donkey serum for one hour at room temperature. Cells were washed three times with PBS and mounted in mounting media with DAPI.

### Immunofluorescence + Telomere FISH

For IF+FISH, cells were grown on acid-washed coverslips. IF was performed as above, with the modification that after permeabilization coverslips were treated with 100 µg/mL RNase A in PBS for 20-30 min at 37 °C followed by thorough rinsing with PBS. After the wash steps following the secondary antibody incubation, coverslips were re-fixed for 15 minutes in freshly made 4% paraformaldehyde in PBS at room temperature, then rinsed well with PBS and dehydrated. The TelC PNA FISH probe (Panagene) was diluted to 200 nM in hybridization buffer (70% deionized formamide, 10 mM Tris (pH 7.4), 0.5% 10x Roche blocking solution) and denatured at 90 °C for 5 minutes. Meanwhile, coverslips were denatured at 85 °C for 5 minutes. Coverslips were inverted onto 75 µL droplets of diluted probe and heated to 85 °C for an additional 5 minutes. Hybridization proceeded overnight at room temperature. Coverslips were then washed and mounted with Prolong Diamond mounting media with DAPI.

### Microscopy

All images were captured on a Zeiss LSM 780 confocal microscope using a 63x oil immersion objective. Zeiss Zen Black software was used for microscope control. Images were cropped, adjusted for brightness and contrast, and pseudo-colored using ImageJ. No non-linear adjustments were used.

### Quantification of colocalized foci

For quantification of colocalized foci, a KNIME workflow was used leveraging spot detection based on wavelet transform^38^. This workflow is available upon request.

### ATRX ChIP-Seq

ATRX ChIP-Seq was performed as previously described^39^, using rabbit anti-ATRX antibody (Santa Cruz, H-300) or a rabbit IgG control. Chromatin was prepared from G292-iDAXX cells plus or minus DAXX induction for two days. After reversing crosslinks, libraries were prepared and sequenced on an Illumina NextSeq 500. Telomere and GC content of reads were determined using TelSeq.^40^

### H3.3 ChIP

ChIP against histone H3.3 was performed using the ChIP-IT kit (Active Motif) as per the manufacturer’s instructions, using rabbit anti-H3.3 polyclonal antibody (GeneTex, GTX112955). After reversing crosslinks, products were blotted onto positively charged nylon membrane (Roche) and crosslinked. Hybridization and visualization steps were performed as per C-circle blots. Images were quantified using ImageJ.

## Acknowledgements

This research was supported by the Intramural Research Program of the NIH, NCI, Center for Cancer Research. We thank the CCR Sequencing core for Sanger sequencing of expression vectors. Microscopy was performed at the CCR Microscopy Core. We thank Eros Lazzerini Denchi for helpful feedback and discussion.

## Author Contributions

S.F.C.S. and P.S.M. conceived the project. S.F.C.S., R.L.W., and M.A.P performed experiments. S.F.C.S., Y.J.Z, and J.L.T.D. analyzed data. S.F.C.S. and P.S.M. wrote the manuscript with input from all authors.

## Data availability

ATRX ChIP-seq data are available upon request.

