## Supplemental Information for "Cancer-associated DAXX mutations reveal a critical role for ATRX localization in ALT suppression"

| | F | R |
|---|---|---|
| L130R | CAATGAGCGCTGCACTGTTCTCAAGGCC | GTGCAGCGCTCATTGATGTAGACATAGAGC |
| L134P | CACTGTTCCCAAGGCCCACTCAGCCAA | GCCTTGGGAACAGTGCAGAGCTCATTGA |
| N153S | CACCTCCAGTGAGCCCTCTGGGAATAACCCTC | GGCTCACTGGAGGTGGTGGCGGCAG |
| S220P | ACCCAGACCCCGCATACCTGCAGGAGGC | ATGCGGGGTCTGGGTCATCCAATTCTGAGAG |
| Y222E | CTCCGCAGAACTGCAGGAGGCACGGTTG | TGCAGTTCTGCGGAGTCTGGGTCATCC |
| E225A | CCTGCAGGCGGCACGGTTGAAGCGTAAGC | CGTGCCGCCTGCAGGTATGCGGAGTCTGG |
| N268S | AGAGGTTAGCAGGCGCATTGAGCGGCTC | CGCCTGCTAACCTCTGGGTAGCGGGTGC |
| A297P | AGAAGGCACCTGCCCGACACAGCCTTG | GGGCAGGTGCCTTCTCTACAGCCCG |
| G321D | AGATGTGGACATCAGGTTACAGGAGCGACG | CTGATGTCCACATCTCGGAAGGCATCC |
| Q469H | CTGGAACACATGCAGGAGGGTCAGGAGG | CTGCATGTGTTCCAGATCCTCCTCCTC |
| D738V | GCTCTCAGTCTCTGATGACCCAGCTTTCTTGTAC | TCAGAGACTGAGAGCACGATGATCTCTTCTGG |

Supplementary Table 1. DNA Oligonucleotides used to generate lentivectors for this study

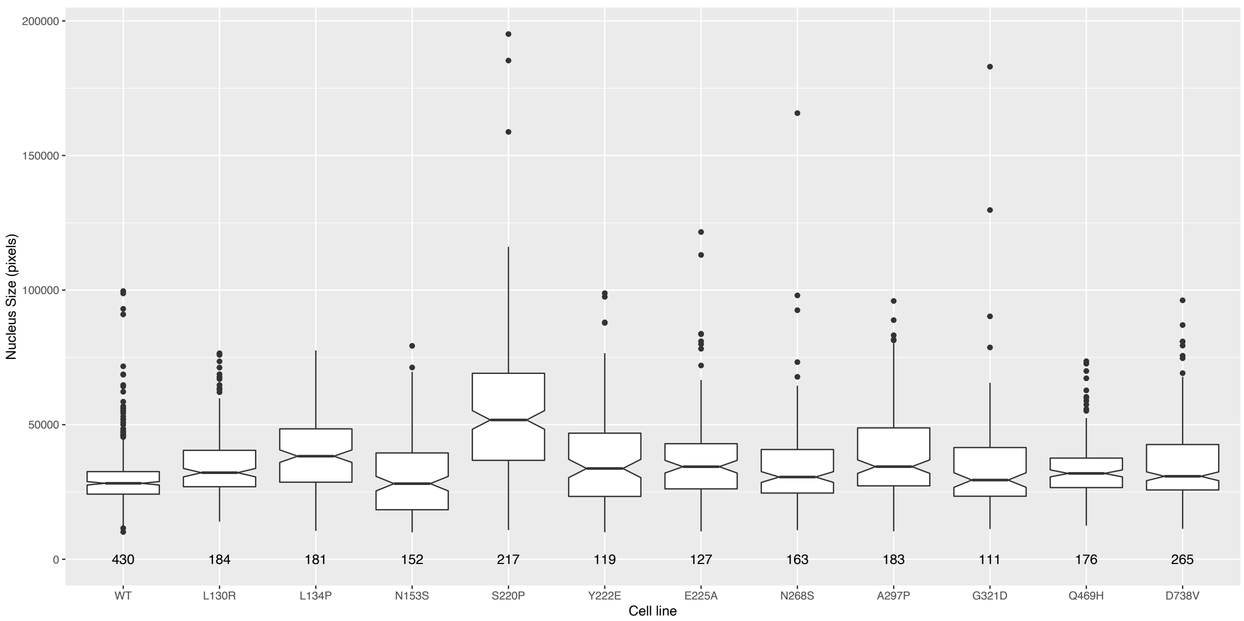


Supplementary Figure 1. Sizes of nuclei in DAXX variant stable lines. Larger nuclei in the S220P cell line suggest a genome duplication event.

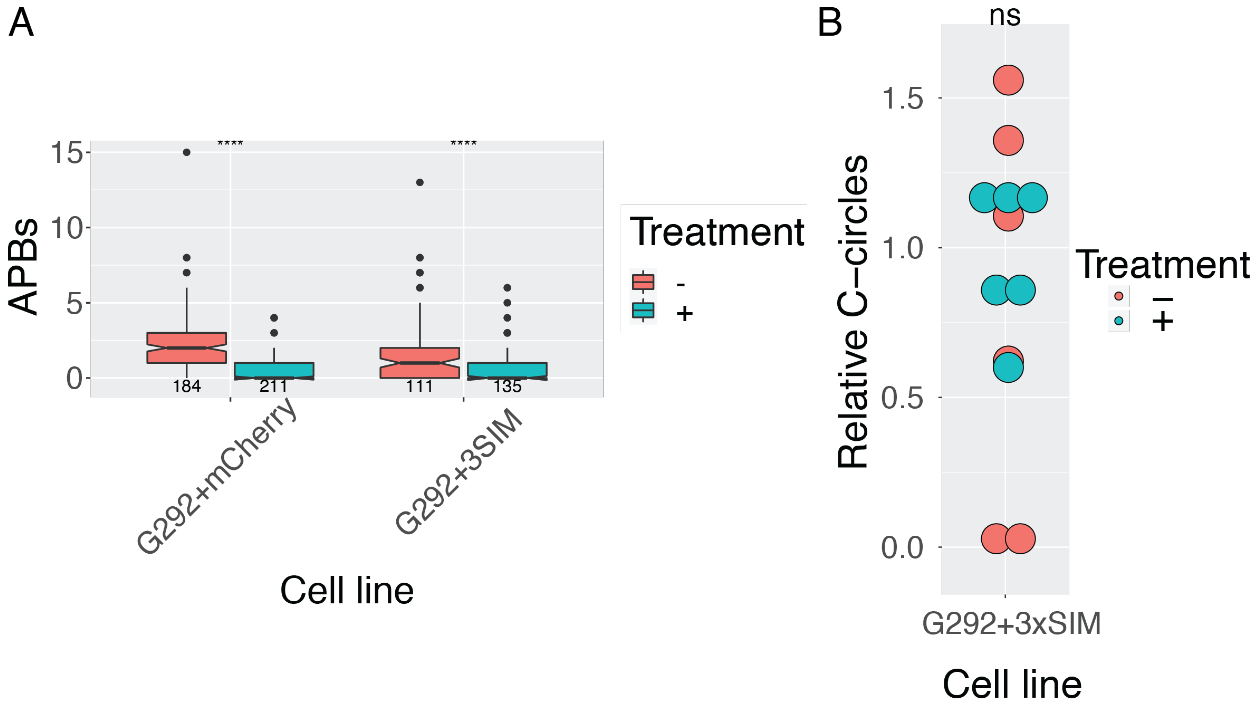


Supplementary Figure 2. APBs are sensitive to induction with doxycycline, but C-circles are not. A. Induction of expression of either mCherry or mCherry-3xSIM with doxycycline decreases APBs per cell significantly. B. Induction of mCherry-3xSIM does not change C-circle levels, implying the reduction in number of APBs does not reflect a *bona fide* change in ALT status.

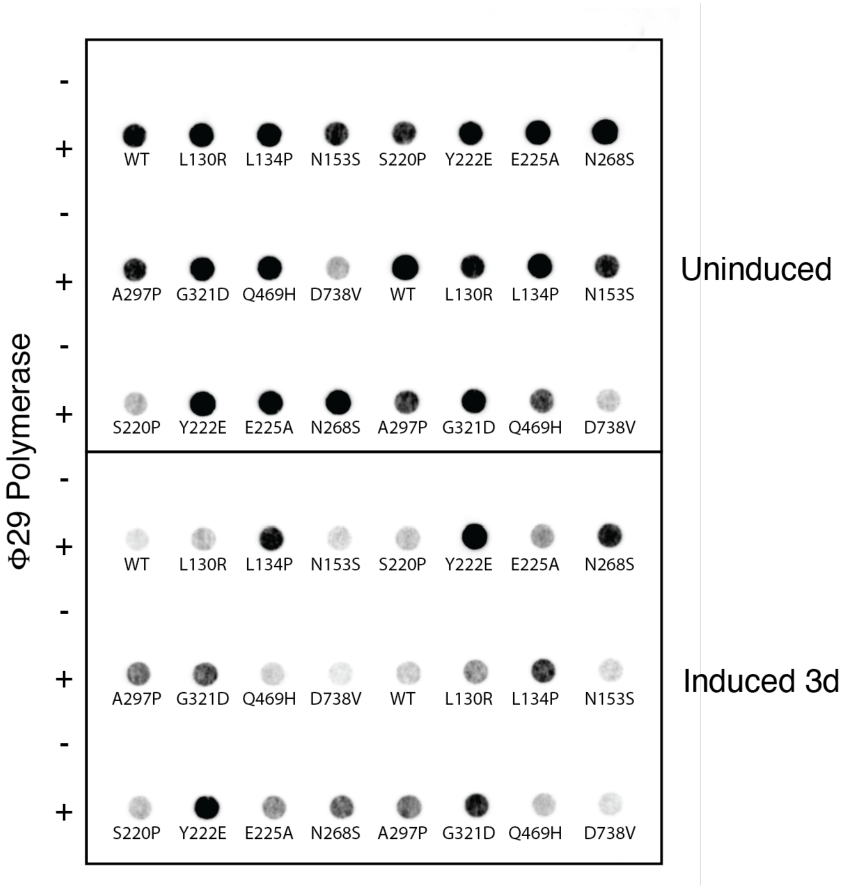


Supplementary Figure 3. Representative C-circle dot blot. Spot intensities were measured using ImageJ, and normalized to the median per-blot intensity for comparison across experiments.
